# *Enterococcus faecalis* manganese exporter MntE alleviates manganese toxicity and is required for mouse gastrointestinal colonization

**DOI:** 10.1101/2020.02.06.936633

**Authors:** Ling Ning Lam, Jun Jie Wong, Kelvin Kian Long Chong, Kimberly A. Kline

## Abstract

Bacterial pathogens encounter a variety of nutritional environments in the human host, including nutrient metal restriction and overload. Uptake of manganese (Mn) is essential for *Enterococcus faecalis* growth and virulence; however, it is not known how this organism prevents Mn toxicity. In this study, we examine the role of the highly conserved MntE transporter in *E. faecalis* Mn homeostasis and virulence. We show that inactivation of *mntE* results in growth restriction in the presence of excess Mn, but not other metals, demonstrating its specific role in Mn detoxification. Upon growth in the presence of excess Mn, an *mntE* mutant accumulates intracellular Mn, iron (Fe), and magnesium (Mg), supporting a role for MntE in Mn and Fe export, and a role for Mg in offsetting Mn toxicity. Growth of the *mntE* mutant in excess Fe also results in increased levels of intracellular Fe, but not Mn or Mg, providing further support for MntE in Fe efflux. Inactivation of *mntE* in the presence of excess iron also results in the upregulation of glycerol catabolic genes and enhanced biofilm growth, and addition of glycerol is sufficient to augment biofilm growth for both the *mntE* mutant and its wild type parental strain, demonstrating that glycerol availability significantly enhances biofilm formation. Finally, we show that *mntE* contributes to infection of the antibiotic-treated mouse gastrointestinal (GI) tract, suggesting that *E. faecalis* encounters excess Mn in this niche. Collectively, these findings demonstrate that the manganese exporter MntE plays a crucial role in *E. faecalis* metal homeostasis and virulence.

## Introduction

Manganese (Mn) is an essential metal for most bacteria and serves as a cofactor for proteins involved in metabolism, DNA replication, respiration, and oxidative stress (1). Accordingly, Mn acquisition contributes to bacterial virulence in multiple bacterial species (2-4). In order to limit bacterial growth and virulence, the host sequesters Mn as a defense response termed nutritional immunity (1, 4-6). To counteract these host-mediated defences, many bacterial pathogens including Enterococci encode dedicated systems to acquire Mn.

Bacteria encode conserved ABC and NRAMP-family transporters for manganese uptake (1, 2). In *Enterococcus faecalis*, three manganese uptake systems have been described: the ABC-type transporter encoded by *efaCBA* and two NRAMP transporters encoded by *mntH1* and *mntH2* (7). These three Mn transport systems are functionally redundant since deletion of all three transporter systems (*efaCBA, mntH1, mntH2*) is required to abrogate intracellular Mn accumulation, rendering the triple mutant severely impaired in growth (7). Furthermore, deletion of both *efaCBA* and *mntH2*, or all three transporters together results in attenuated colonization in rabbit endocarditis and mouse catheter-associated urinary tract infection (CAUTI) models (7). Together these observations demonstrate that the ability to acquire Mn is essential for *E. faecalis* virulence.

In contrast to Mn influx mechanisms that have been characterized in *E. faecalis*, Mn export pathways have not been described. The contribution of Mn export to bacterial pathogenesis and virulence has been demonstrated for some bacterial species (8-15) and is dependent on two widely characterized classes of exporters – MntE, a cation diffusion facilitator (CDF) family protein, and MntP, a 6 transmembrane helix protein typical of LysE transporter family members (8). In *Escherichia coli* and *Neisseria meningitidis*, deletion of *mntP* and *mntX* Mn exporters, respectively, both of which belong to the MntP class of exporters, results in elevated intracellular Mn and increased sensitivity to Mn toxicity (16-18). In the case of *N. meningitidis*, loss of MntX results in reduced bacterial titers recovered from the serum in a mouse sepsis model (17). The CDF family of proteins has been characterized in several bacterial species where they display broad metal specificity (19-23). In *Streptococcus pneumoniae* and *Streptococcus pyogenes*, deletion of the *mntE* results in increased sensitivity to Mn toxicity and intracellular Mn accumulation (9, 10, 24). Similarly, in *Streptococcus suis*, deletion of *mntE* results in increased sensitivity to Mn toxicity and, like *S. pyogenes*, displays increased sensitivity to oxidative stress (10, 12). Additionally, MntE mutants in *S. pneumoniae* and *S. suis* are attenuated in mouse models of infection (9, 12). In *Staphylococcus aureus*, an *mntE* mutant displays increased sensitivity to Mn and oxidative stress when grown in Mn enriched media, and accumulates intracellular Mn (25). Furthermore, loss of *S. aureus mntE* results in reduced mortality in mice after retro-orbital infection (25). Taken together, these reports indicate that Mn import, export, and homeostasis are important for virulence in many pathogens.

In a prior study, we discovered that a putative cation efflux transporter *OG1RF_10589* (*OG1RF_RS03085*) contributed to *E. faecalis* biofilm formation specifically in iron-supplemented media (26). Previous reports showed that *E. faecalis OG1RF_10589* (identified as *czcD* in those studies) transcription was down-regulated when grown in blood (27), and subsequently induced when *E. faecalis* was grown planktonically in both iron (Fe)-supplemented (28) and Mn-supplemented media (29). The goal of this study was to characterize the function of the predicted cation transporter *OG1RF_10589* in biofilm formation. We show that this protein functions in Mn efflux and hence rename the gene *mntE*. An *E. faecalis mntE* mutant grown in excess Mn accumulates intracellular Mn and Fe, but is selectively sensitive only to Mn and not Fe toxicity. However, when *E. faecalis* biofilms were grown in Fe-supplemented media, the conditions in which *mntE* contributed to augmented biofilm formation, only three genes were differentially regulated in the *mntE* mutant compared to wild type: the glycerol catabolic genes were all upregulated (*glpF2, glpK, glpO*). Since we show that glycerol supplementation also promotes biofilm growth, this result suggests that upregulation of glycerol catabolic genes likely contributes to enhanced biofilm growth of the *mntE* mutant in iron-supplemented media that we reported previously. Finally, we demonstrate MntE contributes to colonization of the mouse gastrointestinal (GI) tract, suggesting that maintaining MntE-mediated metal homeostasis confers a fitness advantage to *E. faecalis* in the mammalian host.

## Results

### *mntE* is required for planktonic growth and biofilm formation when Mn is in excess

The *Enterococcus faecalis* OG1RF genome encodes a putative cation efflux transporter (*OG1RF_10589*) and this translated gene product displays 69% and 80% amino acid similarity to the zinc exporter CzcD (Accession no. CWI93218.1) and the manganese exporter MntE (*spy1552*) in *Streptococcus pneumoniae* (9, 10), respectively (see Materials and Methods). MntE belongs to the cation diffusion facilitator (CDF) family of metal efflux pumps (30). The predicted *E. faecalis* cation efflux transporter *OG1RF_10589* also shares 25% amino acid identity to the Mn exporter MneP (formerly called YdfM) in *B. subtilis* and 25% identity to FieF in *E. coli* which, export Mn and Fe, respectively (22, 31). Pairwise alignment of *E. faecalis* MntE showed higher similarity to *S. pneumoniae* and *E. coli* as compared to *B. subtilis* **(Figure S1)**. *E. faecalis OG1RF_10589* possesses a DXXXD motif in transmembrane domain 5 (starting from amino acid 171 in the predicted OG1RF protein) **(Figure S2)** which is typical of CDF transporters (32). CDF transporters possess six putative transmembrane (TM) domains, a signature N-terminal amino acid sequence and a characteristic C-terminal cation efflux domain (33, 34). The experiments below establish *OG1RF_10589* to share properties with MntE and function in Mn homeostasis, so we refer to *OG1RF_10589* as *mntE* henceforth. Based on the amino acid identity between *E. faecalis OG1RF_10589* and Mn transporters in other bacterial species, as well as transcriptional induction of the gene in the presence of both Mn and Fe (28, 29), we hypothesized that *E. faecalis* OG1RF_10589 (MntE) exports Mn and Fe, and that absence of this gene would result in increased sensitivity to metal toxicity and decreased growth in the presence of excess Mn and/or Fe. To test this prediction, we performed planktonic growth assays comparing wild type *E. faecalis* OG1RF to an isogenic *mntE∷Tn* mutant in increasing concentrations of Mn, Fe, zinc (Zn), copper (Cu), and magnesium (Mg). We observed a dose-dependent reduction in growth of the *mntE∷Tn* mutant when grown in Mn supplemented media after 8 hrs of incubation as measured by optical density **(Figure 1A)**. However, there was no growth defect for the *mntE∷Tn* mutant when the media was supplemented with any other cationic metal **(Figure S3)**. Similarly, we observed significantly fewer colony forming units (CFU) (approximately 2-3 log reduction) of *mntE∷Tn* after 6 hours for all Mn concentrations tested **(Figure 1B)**. Complementing the *mntE∷Tn* mutant with *mntE* under control of a nisin-inducible promoter on a plasmid rescued Mn-mediated growth inhibition and restored CFU to near wild type levels after 8hrs of exposure to 2 mM Mn **(Figure 1C-D)**.

**Figure 1.**
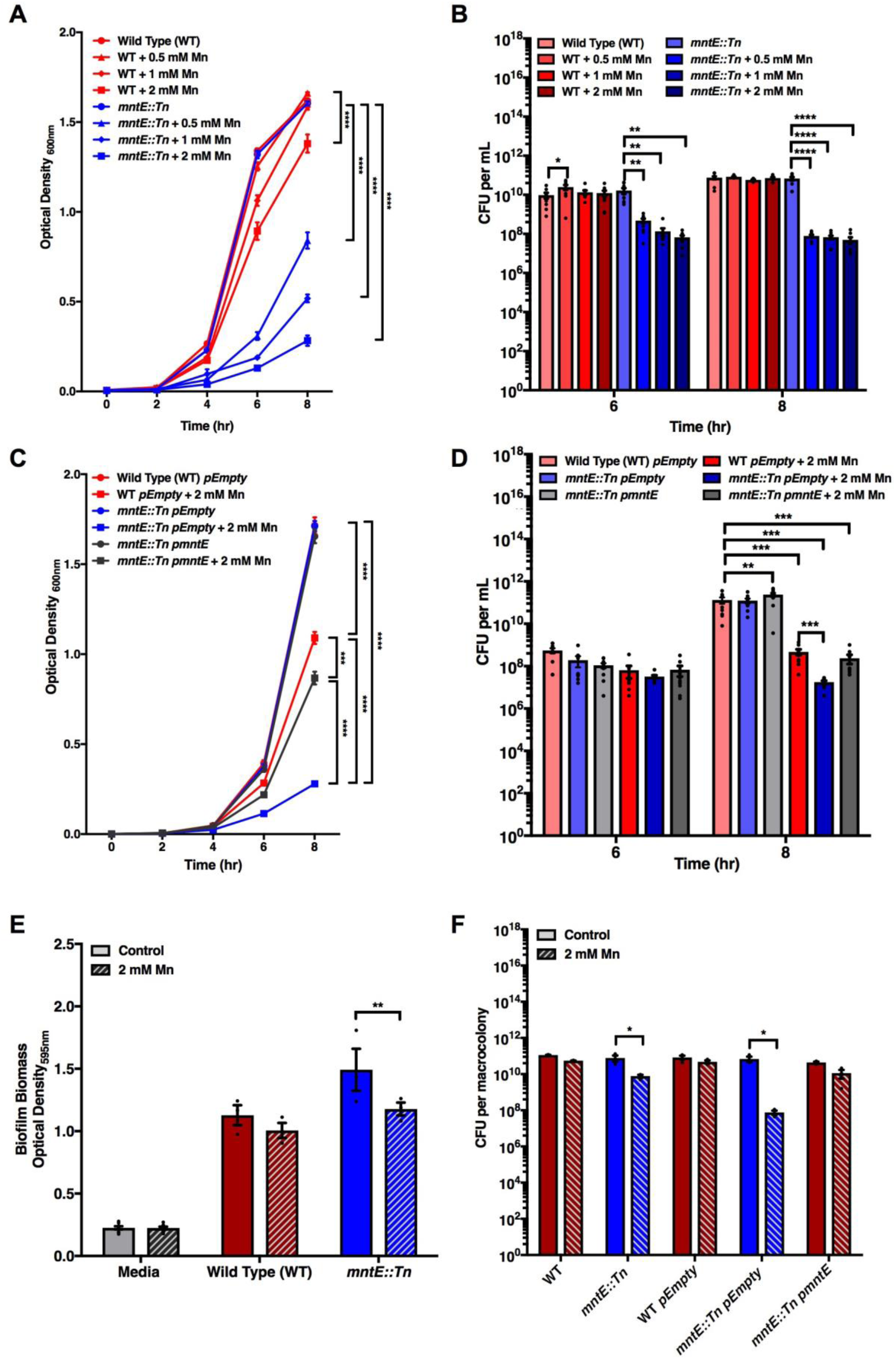
MntE is essential for planktonic and biofilm growth. **(A-D)** Optical density measurement of planktonic growth in Mn supplemented media over the course of 8 hrs (**A**,**C**) and the corresponding CFU enumeration at 6 hrs and 8 hrs time (**B**,**D**). For optical density measurement, statistical analysis was performed at the 8 hr time point using one-way ANOVA with Bonferroni multiple comparison test. For CFU enumerated, statistical analysis was performed using two-way ANOVA with Bonferroni multiple comparison test. Data points represent *n* = 3 experiments, with three independent biological replicates averaged in each experiment. **(E)** *E. faecalis* biofilm biomass quantification grown for 120 hours using crystal violet staining. **(F)** CFU enumeration of macrocolony biofilms. For both biofilm biomass quantification and macrocolony enumeration, statistical analysis was performed using two-way ANOVA with Bonferroni multiple comparison test. Data points represent at least *n* = 3 experiments, with three independent biological replicates averaged in each experiment. Error bar represents standard error of the mean (SEM).

Since the absence of *mntE* leads to Mn-mediated growth inhibition under planktonic conditions **(Figure 1A)**, we next tested whether this was the case for biofilm formation. To address this, we performed static *in vitro* crystal violet (CV) biofilm assays and macrocolony biofilm assays. Using this biofilm accumulation assay, wild type *E. faecalis* biofilm biomass was not significantly altered in the presence of 2 mM Mn. By contrast, the *mntE∷Tn* mutant was attenuated for biomass accumulation **(Figure 1E)**. Similarly, in biofilm macrocolony assays, biofilm CFU were not affected when wild type *E. faecalis* biofilm biomass was grown in 2 mM Mn, but the *mntE∷Tn* mutant had significantly fewer biofilm-associated CFU when grown in excess Mn **(Figure 1F)**. Complementation of *mntE∷Tn* with *mntE in trans* restored biofilm CFU to wild type levels. These results demonstrate that the absence of *mntE* leads to increased sensitivity to Mn during both planktonic and biofilm growth.

### Absence of *mntE* results in intracellular metal accumulation

The ability to regulate intracellular Mn is a key determinant for cell survival and growth. Based on its predicted function in Mn export, we hypothesized that the absence of *mntE* would lead to increased intracellular Mn. To test this hypothesis, we performed inductively coupled plasma mass spectrometry (ICP-MS) on cells isolated from static 24 hr biofilms grown in 2 mM Mn-supplemented media. While we did not observe differences in intracellular metal accumulation when the *mntE∷Tn* mutant was grown in control media **(Figure 2A)**, we observed that wild type *E. faecalis* accumulated more intracellular Mn when grown in Mn-supplemented media and the *mntE∷Tn* mutant accumulated significantly more intracellular Mn compared to wild type, when both were grown in 2 mM Mn-supplemented media **(Figure 2B)**. Complementing the *mntE∷Tn* mutant with *mntE* restored intracellular Mn levels of the *mntE∷Tn* mutant to that of wild type empty vector control strain **(Figure 2B)**. Notably, growth of the *mntE∷Tn* mutant in Mn-supplemented media also resulted in 10-fold more intracellular Mg and 30-fold more intracellular Fe as compared to the wild type strain **(Figure 2B)**. We previously showed that the absence of *mntE* resulted in enhanced biofilm growth in iron supplemented media (26). If MntE also exports Fe, as the data in Figure 2B suggest, the absence of *mntE* should give rise to increased intracellular Fe due to intracellular accumulation. Indeed, we observed that the *mntE∷Tn* grown in Fe-supplemented media accumulated significantly more intracellular Fe as compared to wild type, whereas intracellular Mn and Mg were unchanged compared to wild type **(Figure 2C)**. Complementing the mutant strain with *mntE* resulted in restoration of intracellular Fe to levels observed in wild type empty vector control strain **(Figure 2C)**. These findings suggest that MntE has the capacity to export both Mn and Fe.

**Figure 2.**
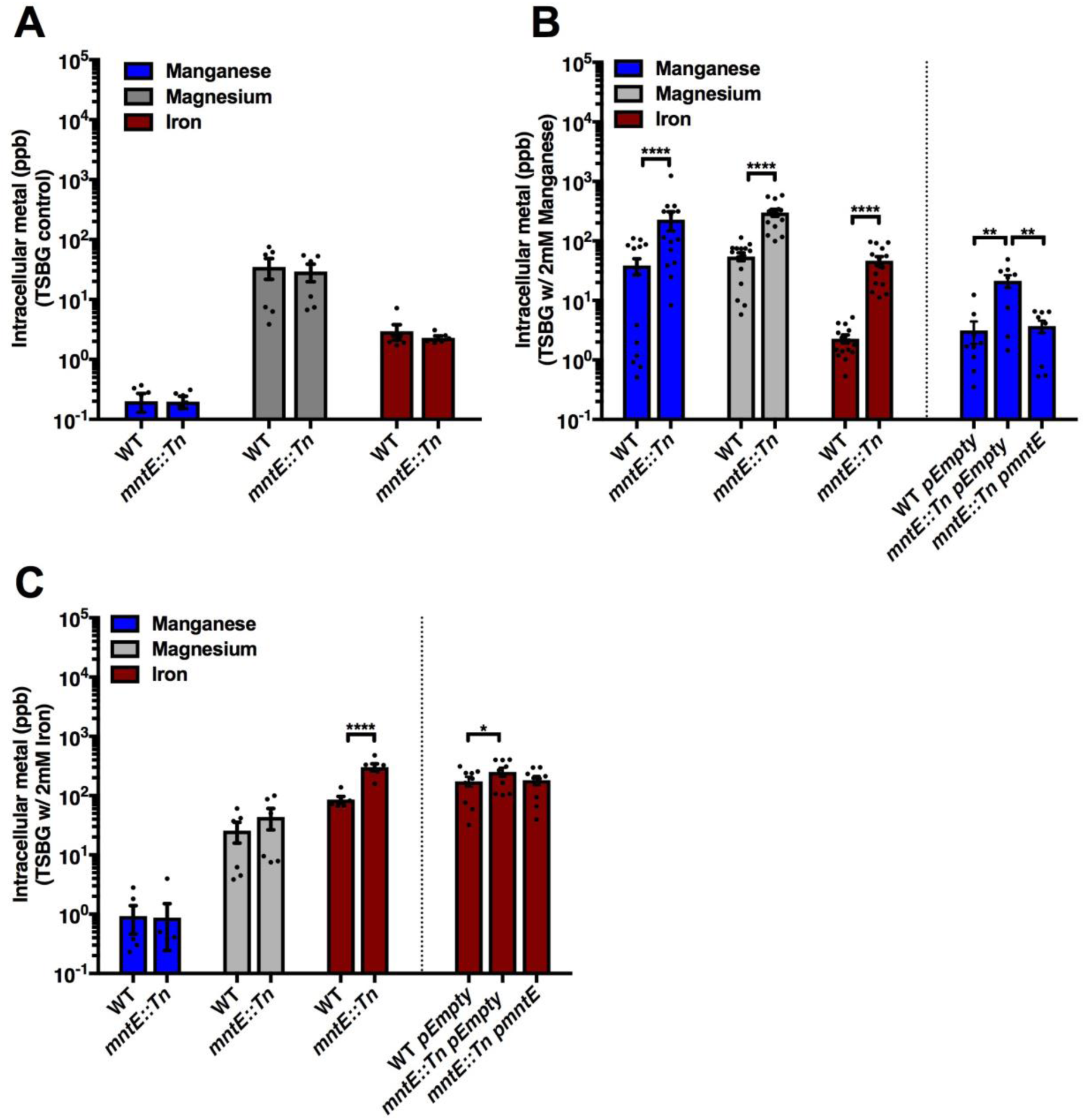
Intracellular manganese, iron, and magnesium content in *E. faecalis* biofilm. ICP-MS analysis of intracellular metals of **(A)** 24hr biofilms grown in control media, **(B)** 2 mM Mn supplemented media and **(C)** 2 mM Fe supplemented media. Data points represent nine independent biological replicates assessed in *n* = 3 experiments. Statistical analysis was performed using two-way ANOVA with Bonferroni multiple comparison test. Error bar represents SEM.

### Magnesium supplementation alleviates manganese-mediated growth inhibition

Since we observed increased intracellular Mg in the *mntE∷Tn* biofilms grown in Mn-supplemented media **(Figure 2)**, we reasoned that increasing intracellular Mg may be a bacterial response to counter accumulated intracellular Mn mediated toxicity as reported in *Bradyrhizobium japonicum* (35). Therefore, we tested if supplemention of Mg would restore growth attenuation of *mntE∷Tn* when grown in 2 mM Mn-supplemented media during planktonic and biofilm growth. Indeed, addition of Mg to Mn-supplemented media restored growth to the *mntE∷Tn* mutant and promoted growth of wild type *E. faecalis* in a dose-dependent manner **(Figure 3A)**. However, in the biofilm assay, we observed significantly attenuated biofilm formation with increasing Mg supplementation for wild type, whereas biofilm of the *mntE∷Tn* mutant was augmented with Mg supplementation **(Figure 3B)**, as it was during planktonic growth **(Figure 3A)**. In the macrocolony assay, the one-log reduction in CFU observed for the *mntE∷Tn* mutant in 2mM Mn-supplemented media was similarly restored at all concentrations of Mg tested **(Figure 3C)**. Alleviation of Mn-mediated growth inhibition of *mntE∷Tn* was specific to Mg, since Fe supplementation did not restore growth (**Figure S4**). Furthermore, 2 mM Mg addition to Mn-supplemented media resulted in reduced intracellular Mn for both wild type *E. faecalis* (3.33-fold) and *mntE∷Tn* (2.21-fold) **(Figure 3D)**. By contrast, supplementing 0.5 mM Mg to Mn-supplemented media resulted in 2-fold and 4-fold increased intracellular Mg in the wild type and *mntE∷Tn* mutant, respectively. Further Mg supplementation significantly reduced intracellular Mg in both wild type and the *mntE∷Tn* mutant **(Figure 3E)**. Together, these observations suggest that Mg supplementation rescues Mn-mediated toxicity and growth inhibition in *E. faecalis*, and that Mg accumulation can impact intracellular Mn pools and modulate biofilm growth.

**Figure 3.**
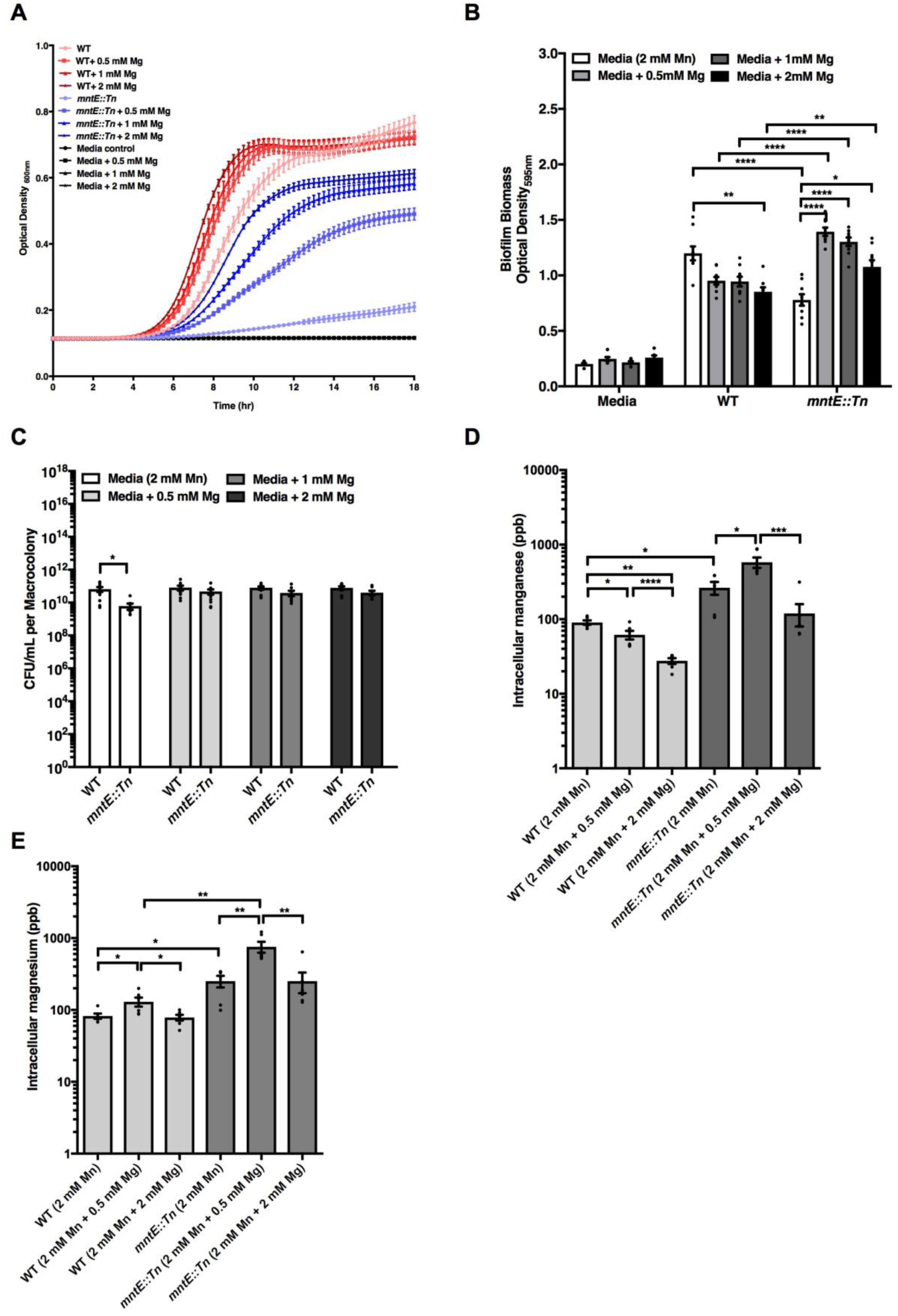
Magnesium supplementation rescue manganese mediated growth inhibition. **(A)** Planktonic growth kinetics. Data points represent *n* = 3 experiments, with three independent biological replicates averaged in each experiment. **(B)** *E. faecalis* biofilm biomass quantification grown for 120 hours using crystal violet staining. **(C)** CFU enumeration of macrocolony biofilms. For both biofilm biomass quantification and macrocolony enumeration, data points represent nine independent biological replicates assessed in *n* = 3 experiments. For both biofilm biomass quantification and macrocolony CFU enumeration, statistical analysis was performed using unpaired t-test with welch’s correction and two-way ANOVA with Bonferroni multiple comparison test respectively. **(D)** ICP-MS analysis of intracellular manganese, **(E)** ICP-MS analysis of intracellular magnesium. For ICP-MS analysis of intracellular manganese and magnesium, data points represent six independent biological replicates assessed in *n* = 2 experiments. Statistical analysis was performed using one-way ANOVA with Bonferroni multiple comparison test. Error bar represents SEM.

### *mntE* expression is manganese responsive in *E. faecalis* biofilm

Since complementation of *mntE* alleviates intracellular Mn accumulation in the *mntE∷Tn* mutant, we hypothesized that *mntE* would be transcriptionally upregulated upon *E. faecalis* biofilm growth in Mn-supplemented media, as previously described for planktonically grown *E. faecalis* (29). We performed qRT-PCR to analyze *mntE* transcript levels and observed a significant increase in expression for wild type *E. faecalis* biofilms grown in Mn-supplemented media compared to normal growth media **(Figure 4)**. Since Mn exposure resulted in upregulation of the *S. pneumoniae* MntE exporter and pilus expression (9), and since pilus expression is critical for *E. faecalis* biofilm formation (36, 37), we hypothesized that pilus expression might be Mn-responsive in *E. faecalis* biofilms as well. Indeed, we observed that *ebpC*, encoding the major subunit of the *E. faecalis* endocarditis and biofilm-associated pilus (Ebp) (36) was also significantly induced in Mn-supplemented media **(Figure 4)**.

**Figure 4.**
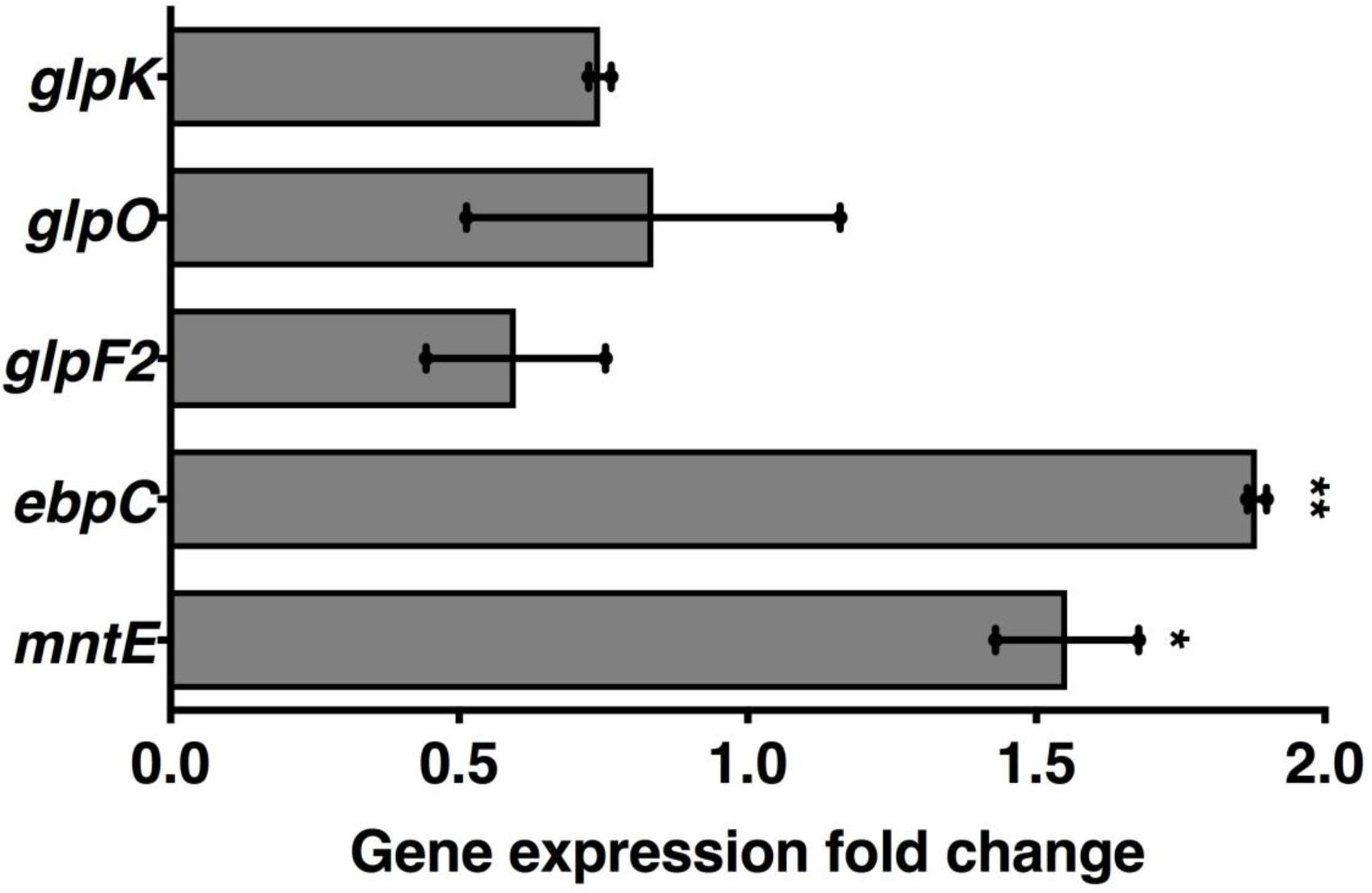
*mntE* expression is induced in *E. faecalis* biofilm upon manganese supplementation. qRT-PCR of *E. faecalis* OG1RF biofilm grown in 2 mM manganese. Data points represents *n* = 2 experiments, with three independent biological replicates averaged in each experiment. Statistical analysis is performed using one-way ANOVA with Fisher LSD test. Error bar represents SEM.

Although the *mntE∷Tn* mutant accumulates intracellular Fe, and *mntE* is induced in response to Fe during planktonic growth (28), *mntE* is not upregulated in *E. faecalis* biofilms when grown in Fe-supplemented media (data not shown). However, we previously showed that the absence of *mntE* resulted in augmented biofilm formation in Fe-supplemented media (26). To identify other Fe-regulated genes that might contribute to Fe-augmented biofilm formation, we performed RNA sequencing and compared transcriptional profiles of wild type and *mntE∷Tn E. faecalis* biofilms grown in Fe-supplemented media. Strikingly, the only differentially regulated genes were the upregulation of glycerol catabolic genes (*glpF2, glpO, glpK*) in the *mntE* mutant in response to Fe when compared to the non-iron supplemented media TSBG control **(Table S1)**. We speculated that glycerol serves as an energy source to promote biofilm growth for the *mntE∷Tn* mutant in Fe-supplemented media. We were unable to simultaneously delete both *mntE* and *glpF2* in order to test this hypothesis. Instead, increasing glycerol concentrations in the growth media enhanced biofilm formation in both the wild type control and *mntE∷Tn* mutant, regardless of Fe supplementation (**Figure S5**). By contrast, these three glycerol catabolic genes (*glpF2, glpO, glpK*) were not upregulated under Mn supplemented conditions in wild type *E. faecalis* biofilm **(Figure 4)** and global transcriptional analysis showed that these genes are not Fe responsive in wild type OG1RF biofilm grown in Fe-supplemented media (38). Taken together, these results indicate that upregulation of glycerol catabolic genes is specifically observed in the absence of *mntE* when intracellular Fe levels are high and that glycerol supplementation contributes to biofilm growth.

### Absence of MntE does not alter oxidative stress tolerance in *E. faecalis*

Since the absence of *mntE* results in intracellular Mn accumulation, we speculated that accumulation of Mn may alter *E. faecalis* oxidative stress tolerance. The increased availability of intracellular Mn could enhance Mn-dependent antioxidant defenses, as has been reported in *Streptococcus spp*. (10, 12). Alternatively, increased intracellular Mn could lead to increase sensitivity to oxidative stress as reported for *Xanthomonas oryzae* and *S. pyogenes* (8, 12, 39). However, the *E. faecalis mntE∷Tn* mutant did not display altered sensitivity to oxidizing agents when compared to wild type **(Figure S6A-B)**, nor did hydrogen peroxide production significantly change when compared to wild type, as has been reported for *S. pneumoniae* when Mn is in excess (9) **(Figure S6C)**. Therefore, we conclude that increased intracellular Mn does not impact oxidative stress tolerance in *E. faecalis*.

### MntE is required for *E. faecalis* expansion in the mouse GI tract

Given the importance of Mn acquisition for *E. faecalis* virulence (7) and the role of MntE in Mn homeostasis, we tested whether MntE contributes to *E. faecalis* virulence. Using an antibiotic-treated mouse model of gastrointestinal (GI) tract colonization, we observed that the *mntE∷Tn* mutant was significantly attenuated for colonization in the cecum, small intestine, and colon as compared to wild type *E. faecalis* **(Figure 5)**. These results suggest that the GI tract represents a natural reservoir abundant with Fe, Mn, or other MntE-effluxed metals. Indeed, ICP-MS analysis of GI tissue showed the presence of Mn and Fe **(Figure S7)**. Therefore, our results demonstrate that MntE and Mn homeostasis, and potentially MntE-mediated Fe homeostasis, are important for *E. faecalis* colonization of the antibiotic-treated mouse GI tract.

**Figure 5.**
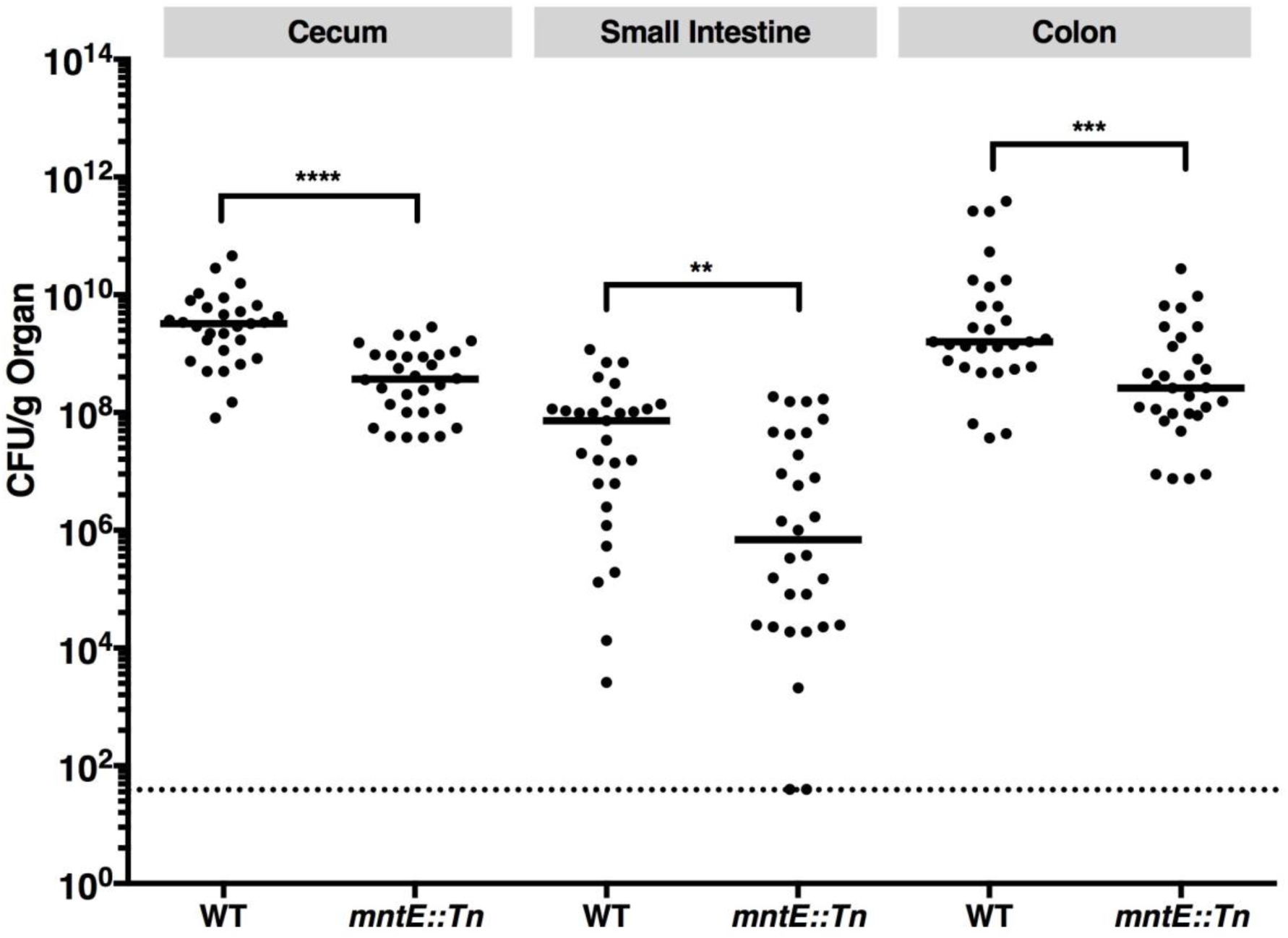
MntE is required for colonization in mouse gastrointestinal tract. Bacterial CFU in GI tissues. Data points represent CFU recovered from each mouse (for WT *n* = 30; for *mntE*∷Tn mutant *n* = 30) collected in three independent experiments. Statistical significance was determined by Mann-Whitney test. Black line indicates the median. Dotted line indicates limitation of detection at CFU of 40.

## Discussion

We previously showed that the absence of MntE resulted in enhanced iron-augmented biofilm (26), however the role of MntE and its contribution to iron-augmented biofilm were not characterized. We show that MntE is essential for Mn homeostasis to prevent Mn toxicity, and contributes to changes in intracellular Fe and Mg pools, which in turn can alter glycerol catabolism and growth. Finally, we demonstrate that MntE is required for *E. faecalis* expansion in the mouse gastrointestinal (GI) tract.

Metal exporters serves to maintain intracellular metal homeostasis. The observation that Mg and Fe accumulate within *E. faecalis mntE* mutant cells during growth in Mn-supplemented media suggests either that MntE can efflux Mg and Fe, or that growth in Mn-supplemented media results in the increased import of these metals. Although Mg^2+^ has similar ionic radii to Fe^2+^ and Mn^2+^, we are not aware of any reports documenting Mg^2+^ efflux from CDF family of proteins (30) or Mg transporters in *E. faecalis*. However, conserved families of bacterial proteins have been identified for Mg uptake and efflux (40). Therefore, these findings together suggest that the observed Mg accumulation in the *E. faecalis mntE* mutant may be mediated by transporters which have yet to be characterized in *E. faecalis*. With regard to Fe accumulation, and consistent with the possibility that MntE effluxes Fe, *mntE* was previously reported to be upregulated when *E. faecalis* was grown in Fe-supplemented media planktonically (28), and the absence of MntE resulted in increased intracellular Fe. While the absence of MntE also similarly resulted in intracellular Mn accumulation, no growth inhibition for the *mntE∷Tn* mutant was observed even at the highest Fe concentration tested as compared to Mn in this study. Based on these data, we speculate that MntE may be the sole exporter for Mn in *E. faecalis*, whereas MntE may be one of several redundant export systems for Fe.

Altered Mn homeostasis affects sensitivity to oxidative stress and has been demonstrated to attenuate virulence in *S. pyogenes* (11, 41), *S. pneumoniae* (9, 42), and *S. aureus* (43). Given the sequence and functional conservation between *E. faecalis* MntE and MntE from other Gram positive species, we examined its role in *E. faecalis* oxidative stress tolerance and virulence. While we found no evidence for a role of *E. faecalis* MntE during growth in the presence of oxidative damaging agents, it contributes to growth in the antibiotic-treated GI tract. A major factor in *E. faecalis* adhesion, virulence, and biofilm formation is the sortase-assembled pilus (Ebp) (37, 44-46). However, transcription of the gene encoding the major pilus subunit *ebpC* is upregulated in Mn-supplemented media, even when biofilm biomass accumulation is attenuated compared to wild type. Since Ebp are important for GI colonization (47), and since the *mntE* mutant is attenuated in mouse GI colonization, we speculate the upregulation of pilus expression observed *in vitro* either does not occur in the mouse GI tract, or that pilus expression *in vivo* is insufficient to complement the virulence defect of the *mntE* mutant.

In this study, we observed that when Mn is in excess, an *E. faecalis mntE* mutant accumulates intracellular Mn, Fe, and Mg. It is likely that altered intracellular metal homeostasis may be the driving force underlying Mn-mediated growth inhibition and the *in vivo* virulence defect. Multiple mismetallation outcomes could be at play resulting in its sensitization to Mn toxicity. The accumulation of intracellular Mg in the absence of *mntE*, coupled with the ability of Mg supplementation to reduce intracellular Mn, restore growth, and protect from Mn toxicity, suggests that mismetallation of Mg-metalloproteins by Mn may be an underlying reason for *E. faealis* growth inhibition. Magnesium can serve as a cofactor for Mg-dependent enzymes, and can help to stabilize protein complexes and cellular structures (48, 49). Due to the similar ionic radii of these divalent ions, we postulate that Mn cations can displace Mg. The displacement of Mg could in turn result in non-functional or altered function of the metalloprotein. Although there is no evidence for this in *E. faecalis* to date, this idea has been proposed in other bacterial spp. In *B. subtilis*, the loss of *mpfA*, encoding a Mg efflux pump, leads to increased intracellular Mg and suppressed Mn toxicity (50). Despite the increased sensitivity to Mg toxicity, the *mpfA* mutant is less sensitive to Mn toxicity. Further, supplementing the growth media with Mg rendered both wild type *B. subtilis* and its Mn efflux mutant (Δ*mneP*Δ*mneS*) less sensitive to Mn intoxication, and also less sensitive to Fe, Co and Zn intoxication (50). In *Bradyrhizobium japonicum*, removal of Mn from Mg-limited media partially restores growth defects due to depletion of Mg, which suggest that the presence of Mn under Mg-limited condition is toxic to *B. japonicum* (35). Consistent with the speculation that Mn and Mg can competitively bind to metalloproteins and alter protein function, supplementing *B. japonicum* with either metal enhances activity of Mg-dependent isocitrate dehydrogenase; by contrast, addition of Mn inhibited Mg-dependent isocitrate lyase (35). Additionally, activity of another *B. japonicum* Mg-dependent enzyme, 5-aminolevulinic acid (ALA) dehydratase, was 3-fold higher with Mn as co-factor as compared to Mg (35). Despite the limited literature describing mismetallation, these findings suggest that mismetallation of Mg-metalloproteins by Mn can alter enzymatic activity and affect growth, and this may be relevant for *E. faecalis*.

We speculate that increased Fe levels observed in the *E. faecalis mntE* mutant may serve to maintain the intracellular Fe/Mn ratio necessary for cellular processes under Mn stress. Altered metal homeostasis occurs when bacteria are under Mn stress, whereby the accumulation of intracellular Mn is accompanied with changes in Fe and Cu levels, as have been described in multiple bacterial species. Previously we have also shown that increased intracellular Fe is accompanied with increased Cu levels when *E. faecalis* biofilm is grown in iron supplemented media (26). In *S. pneumoniae*, deletion of *mntE* results in increased intracellular Mn and similarly, Fe and Cu intracellular levels are increased (24). Growth of the *S. pneumoniae mntE* deletion mutant under Mn stress resulted in upregulation of genes involved in both Fe and Cu uptake (9, 51, 52), and these observations are consistent with the intracellular accumulation of these metals (24). Similarly, in *E. coli*, overexpression of the Mn exporter encoded by *mntS*, or deletion of the Mn exporter encoded by *mntP*, resulted in elevated intracellular Mn (16, 18). However, overexpression of *mntS* resulted in decreased intracellular Fe, due to downregulation of Fur-regulated iron uptake genes (18, 53, 54). In the context of *E. coli*, it was proposed that Mn can substitute for Fe, thus it is likely that Mn-bound Fur is a capable repressor for iron acquisition gene expression. In *S. aureus*, loss of *mntE* expression resulted in elevated intracellular Mn and reduced intracellular Fe (25). It has been suggested that the elevated intracellular Mn drives repression of the PerR regulon which limits oxidative stress responses and Fur-dependent expression of iron acquisition systems in *S. aureus* (55, 56). Altogether these reports demonstrate that bacteria tightly regulate intracellular Mn/Fe ratios, and altered homeostasis of these transition metals can alter gene transcription and growth. Therefore, it is likely that *E. faecalis* employ similar strategies to regulate intracellular Fe/Mn ratios, and alteration of these ratios can impact global gene transcription. We speculate that inactivation of *mntE* did not greatly impact growth in normal media or oxidative stress tolerance due to the presence of redundant antioxidant enzymes in *E. faecalis*. Future studies should focus on the transcriptional changes including *fur* and *perR* regulon under altered intracellular Fe/Mn ratios due to deletion of *mntE*, and how these genes impact intracellular metal homeostasis.

To elucidate mechanisms involved in enhancement of iron-augmented *E. faecalis* biofilm formation by an *mntE* mutant, we discovered that glycerol catabolic genes (*glpF2, glpO, glpK*) were induced in the *mntE∷Tn* mutant when grown in iron-supplemented conditions that also drive intracellular iron accumulation. It is unclear how iron might stimulate glycerol catabolism, but we do know that *E. faecalis* has two glycerol catabolic pathway, one which is dependent on ATP-mediated phosphorylation of glycerol by glycerol kinase (GlpK) to yield glycerol-3-phosphate (glycerol-3-P) (57). Here, we propose a model in which upregulation of the glycerol importer (*glpF2*), alpha-glycerophosphate oxidase (*glpO*), and glycerol kinase (*glpK*) is driven by the presence of increased intracellular Fe. Consistent with this idea, an *E. faecalis* V583 *fur* deletion mutant is incapable of repressing iron uptake, and when grown in iron supplemented media, displayed significantly increased transcription of glycerol dehydrogenase and glycerol kinase (*glpK*) (58). The relationship between glycerol catabolism and Fe availability is unclear at this time. Since Fe may function as a biocatalyst for oxidation of glycerol (59) and is an important transition metal for microbial growth (60), we speculate that Fe positively impacts glycerol uptake and the increased uptake of glycerol, which in turn are converted to glycerol-3-phosphate (G3P) in glycolysis, drives increased energy production and increased biofilm growth.

Collectively, these findings suggest that MntE is a Mn exporter. Since MntE is conserved across a number of gram positive and gram negative bacteria, we propose that this Mn efflux system is a common strategy for Mn homeostasis in bacteria. In *E. faecalis*, we establish that efflux of Mn is vital for growth and successful colonization in the gastrointestinal tract (GI), and that the Mn exporter MntE may be a promising target in developing new therapeutics for patients suffering from VRE dominated intestinal microbiota who are more susceptible to nosocomial infections (61-65).

## Material and methods

### Bacterial Strains and Growth Conditions

*Enterococcus faecalis* was grown in Brain Heart Infusion broth (BHI) and cultured at 37 °C under static or shaking (200rpm) conditions, as indicated below. Preparation of inocula for biofilm and planktonic assays was performed as previously described (26). Bacterial strains used are listed in **Table 2**. Where appropriate, strains harbouring pMSP3535 plasmids were selected using 100 ug/mL erythromycin (Sigma Aldrich, USA) and induction of gene expression was performed using 5 ug/mL nisin from *Lactococcus lactis* (Sigma Aldrich, USA). BHI was supplied by Becton, Dickinson and Company, Franklin Lakes, NJ. TSB and agar was supplied by Oxoid Inc., Ontario, Canada. Metals were filtered sterilized and supplemented during medium preparation in autoclaved TSBG media. For experiments using ferric chloride only, metal is supplemented in TSBG media and autoclaved together. Magnesium chloride anhydrous ≥98%, copper chloride dihydrate ≥99%, ferric chloride anhydrous ≥99% and heme ≥90% were supplied by Sigma Aldrich, St Louis, MO, USA. Manganese chloride tetrahydrate and zinc chloride were supplied by Merck Millipore, Singapore.

### Protein homology determination

*E. faecalis* OG1RF MntE (GenBank: AEA93276.1) amino acid sequence (389 amino acids) was queried against the non-redundant GenBank CDS including *Streptococcus pneumoniae, Bacillus subtilis* and *Escherichia coli* taxonomy using the NCBI blastp online tool.

### General cloning techniques

Nucleotide sequence of *mntE* is obtained from the *E. faecalis* OG1RF genome via BioCyc (66). The Wizard genome DNA purification kit (Promega Corp., Madison, WI) was used for isolation of bacterial genomic DNA (gDNA), and Monarch® Plasmid miniprep Kit (New England BioLabs, Ipswitch, MA) was used for purification of plasmid for gene expression and construction of complement mutant. The Monarch® DNA Gel Extraction Kit (New England BioLabs, Ipswitch, MA) was used to isolate PCR products during PCR. In-Fusion HD Cloning Kit (TaKara Bio, USA) was used for fast, directional cloning of DNA fragments into expression vector. All plasmids used in the study are listed in **Table S2**. T4 DNA ligase and restriction endonucleases were purchased from New England BioLabs (Ipswitch, MA). Colony PCR was performed using Taq DNA polymerase (Thermo Fisher Scientific, Waltham, MA, USA) and PCR of gene of interest for plasmid construction was performed using Phusion DNA polymerase (Thermo Fisher Scientific, Waltham, MA, USA). Ligations were transformed into *E. coli* Dh5α cells. Plasmids derived in this study were confirmed by sequencing of purified plasmid.

### Strain construction

To construct *mntE* complementation plasmid, primers (mntE_F’ and mntE_R’; **Table S3)** were designed with BamHI restriction site and SmaI restriction sites flanking the gene of interest, to generate DNA fragments as templates. In-Fusion cloning (Takara Bio USA Inc.) was performed using primers (mntE_F’_Infusion and mntE_R’_Infusion) with at least 15 bp complementary sequence for ligation into the nisin-inducible vector pMSP3535, also digested with the same restriction enzymes. pMSP3535∷*mntE* plasmid was generated in *E. coli* Dh5α, verified by sequencing, and transformed into *E. faecalis* as described previously (37).

### Biofilm Assay

Bacterial cultures were normalized as previously described (26),inoculated in TSBG in a 96-well flat bottom transparent microtiter plate (Thermo Scientific, Waltman, MA, USA), and incubated at 37°C under static conditions for 5 days unless specified otherwise. Strains harboring pMSP3535 complementation plasmid was grown in the presence of erythromycin. Adherent biofilm biomass was stained using 0.1% w/v crystal violet (Sigma-Aldrich, St Louis, MO, USA) at 4°C for 30 minutes. The microtiter plate was washed twice with PBS followed by crystal violet solubilization with ethanol:acetone (4:1) for 45 minutes at room temperature. Quantification of adherent biofilm biomass was measured by absorbance at OD_595nm_ using a Tecan Infinite 200 PRO spectrophotometer (Tecan Group Ltd., Männedorf, Switzerland).

### Plate-Assisted Planktonic Growth Assay

Bacterial cultures were normalized as previously described (26) and further diluted by a dilution factor of 200. Diluted cultures were then inoculated into fresh media at a ratio of 1:25, which is 8 µL of the inoculum in 200 µL of media, incubated at 37°C for 18 hours, and absorbance at OD_600nm_ was measured using a Tecan Infinite 200 PRO spectrophotometer (Tecan Group Ltd., Männedorf, Switzerland) at 15 minute intervals (with shaking prior to each measurement).

### Planktonic Growth Kinetic Assay

Bacterial cultures were normalized as previously described (26) and inoculated into fresh media at a ratio of 1:1000 in 30 mL of media in 50 mL conical tubes. The tubes were incubated with shaking for 8 hours at 200 rpm, 37°C. At indicated time intervals, 100 μL and 1 mL of culture was removed for colony forming units (CFU) enumeration and optical density measurement, respectively.

### Macrocolony Assay

Bacterial cultures were normalized as previously described (67) and spotted onto TSBG agar plate at 5μL per spot. Agar plates were supplemented with metals where appropriate and incubated for 120 hours at 37°C. Macrocolonies were excised, vortexed and resuspended in 3 mL PBS, and serially diluted for colony forming unit (CFU) enumeration. 5μL of dilution from each well was spotted onto the agar plates and incubated at 37°C overnight for subsequent calculation of CFU/mL.

### Quantitative Real time PCR (qRT-PCR) and RNA sequencing

Biofilms were grown in a 6-well plate for 24 hours at 37°C under static conditions. Post incubation, spent media was removed and biofilms were suspended in PBS prior being dislodged using cell scraper. Biofilm cultures were centrifuged at 14,000 rpm for 2 minutes at room temperature to remove supernatant. Biofilm cell pellet was incubated with lysozyme from chicken egg white (10mg/ml) (Sigma Aldrich, USA) for 30 minutes at 37°C, and centrifuged at 14,000 rpm for 2 minutes at room temperature to remove supernatant prior to cell lysis. RNA extraction was performed in a Purifier® filtered PCR enclosure using the PureLink™ RNA mini kit (Invitrogen, USA) according to the manufacturer’s instructions. RNA purification and removal of DNA was performed using TURBO DNA-free™ kit (Thermo Fisher, USA) and Agencourt® RNAClean® XP Kit (Beckman Coulter, USA). Measurement of RNA yield and quality was performed using Qubit® RNA HS assay kit (Thermo Fisher, USA) and RNA ScreenTape System and 2200 TapeStation (Agilent, USA). Synthesis of cDNA was performed using SuperScript III First-strand (Invitrogen, USA). Quantitative real-time PRC using cDNA was performed using KAPA SYBR fast qPCR master mix kit (Sigma Aldrich, USA) and Applied Biosystems StepOne Plus Real-Time PCR system. The expression of *ebpC, ebpR, mntE* and *gyrA* were measured using primer pairs listed in **Table 3**. For each primer set, a standard curve was established using genomic DNA from *E. faecalis* OG1RF. Normalized amount of cDNA were used to determine relative fold change in gene expression as compared to *E. faecalis* OG1RF biofilm grown in TSBG. For RNA sequencing, ribosomal RNA depletion was performed after RNA purification using Ribo-Zero™ rRNA removal kit (Illumina, USA). cDNA library synthesis was performed using NEBNext RNA First-strand and NEBNext Ultra directional RNA Second-strand synthesis module (New England BioLab, US). Transcriptome library preparation was performed using 300bp paired end illumina sequencing.

### Mouse Gastrointestinal Tract (GI) Colonization Model

Six week old male C57BL/6NTac mice were administered ampicillin (VWR, USA) in their drinking water (1 g/L) for 5 days as previously described (62, 68). Mice were then given one day of recovery from antibiotic treatment prior to administration of approximately 1-5 × 10^8^ CFU/ml *E. faecalis* (OD_600nm_ 0.5) in the drinking water for 3 days as previously described (69). Before and after infection, mice were monitored for signs of disease and weight loss. All animal experiments were approved and performed in compliance with the Nanyang Technological University Institutional Animal Care and Use Committee (IACUC). At the indicated timepoints, the small intestine, colon, and cecum were harvested. Tissue samples were homogenised in PBS, serial diluted in PBS, and spot-plated on BHI agar with 10 mg/L colistin, 10 mg/L nalidixic acid, 100 mg/L rifampicin, 25 mg/L fusidic acid for CFU enumeration. All antibiotics were obtained from Sigma Aldrich, USA.

### Statistical analyses

Data from multiple experiments were pooled, and appropriate statistical tests applied, as indicated in the respective figure legends. Statistical analyses were performed with GraphPad Prism 6 software (GraphPad Software, San Diego, CA). An adjusted P value of <0.05 was considered statistically significant.

## Supplementary methods

are provided for supplementary experiments.

### Statement of Contribution

L.N.L and K.A.K conceived of and designed the study, analyzed the data, and wrote the manuscript. L.N.L performed all of the experiments. J.J.W assisted with animal experiments and K.K.L.C analyzed the RNA sequencing data. All authors edited and approved the final manuscript.

## Acknowledgments

We are grateful to Jenny Dale and Gary Dunny for supplying us with *E. faecalis* OG1RF transposon mutants used in this study. We are also thankful to Jose Lemos for critical review of this manuscript. This work was supported by the National Research Foundation and Ministry of Education Singapore under its Research Centre of Excellence Programme and by the Ministry of Education Singapore under its Tier 2 programme (MOE2014-T2-2-124).

## Supplementary Materials & Methods

### Oxidant Stress Challenge

Bacterial cultures were normalized to OD 0.7 as previously described (67), and added at 1:25 ratio to media. Supplementation of menadione or hydrogen peroxide stimulate oxidative stress. Bacterial cultures were allowed to grow for 2 h at 37°C static condition prior to CFU enumeration.

### Hydrogen Peroxide Quantification

Overnight bacterial cultures were normalized to OD 0.7 as previously described (67), diluted 1:25 in fresh media, and grown for 2 h at 37°C without shaking. After incubation, hydrogen peroxide quantification was performed using ROS-Glo™ H2O2 Assay (Promega, USA) according to manufacturer’s instructions.

### Inductively Coupled Plasma Mass Spectrometry (ICP-MS)

Biofilms are cultured under static condition at 37°C for 24 hrs. After incubation, biofilms are scraped, resuspended in 1mL PBS and normalized to OD 1. Normalized biofilms are pelleted at 14,000 rpm for 2 minutes, and supernatant was discarded. Preparation of cell pellets for ICP-MS was performed as previously described (26) with minor modifications. Cell pellets are suspended in 300 uL of lysozyme from chicken egg white (20 mg/ml) (Sigma Aldrich, USA) (20mg/mL) for 30 minutes at 37°C, washed with 1 mL PBS and pelleted. At a ratio of 2:1, 70% nitric acid (Sigma Aldrich, USA) and 30% hydrogen peroxide (Sigma Aldrich, USA) was added to normalized lysozyme treated biofilm cells and left under room temperature for 3 days to allow complete digestion. The digested samples were diluted with 3.4 mL LC-MS grade water and filtered using 0.2 um membrane, prior to analysis using ICP-MS. Analysis of trace metals in samples were performed using ICP-MS model Elan-DRCe, Meinhard Nebulizer model TR-30-C3 (Perkin Elmer; Model: N8122006 (Elan Standard Torch)).

## Supplementary Figure Legends

**Figure S1.**
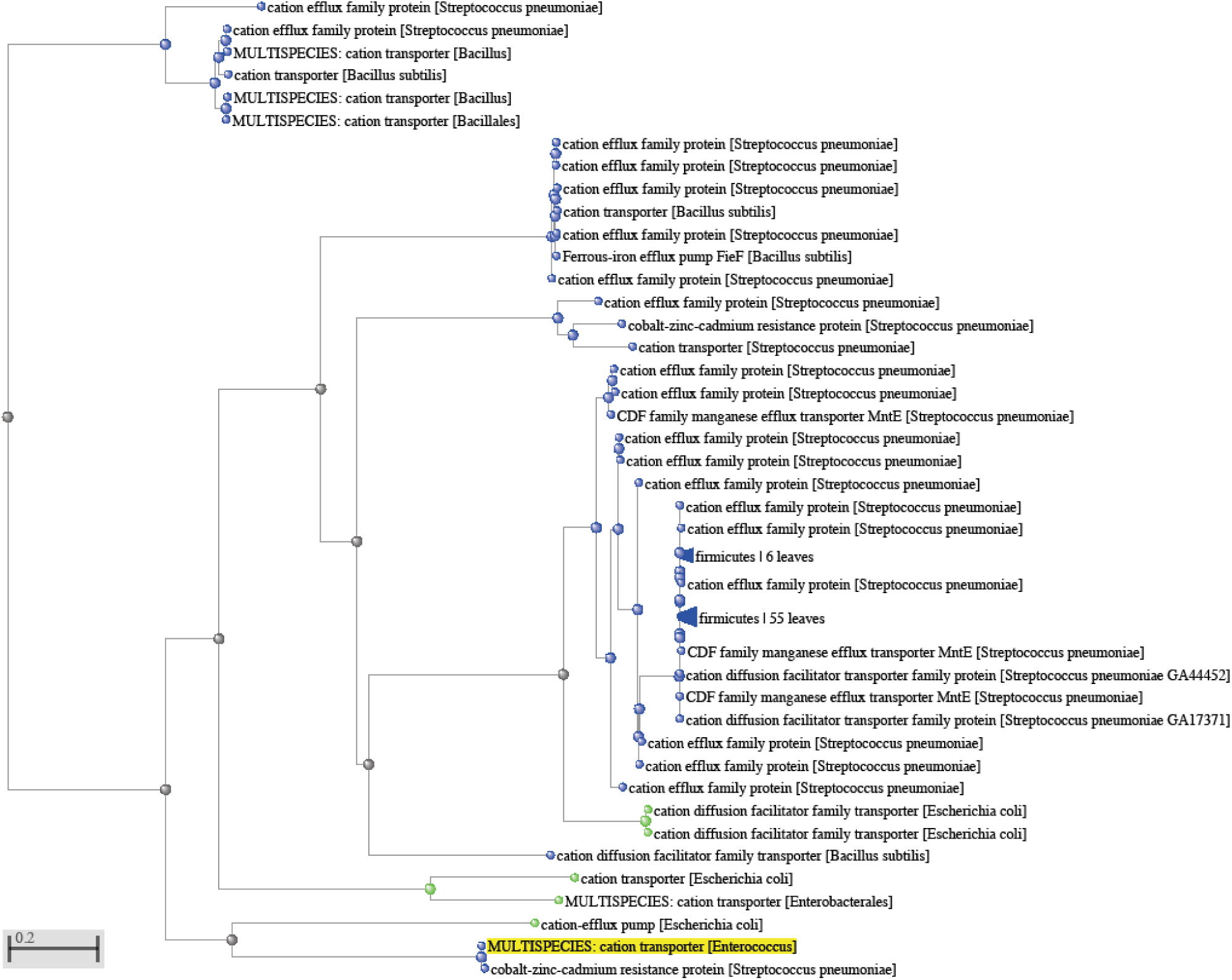
Phylogenetic tree summary of E. faecalis MntE in comparison with *Streptococcus spp*., *Bacillus spp*. and *Escherichia spp*.

**Figure S2.**
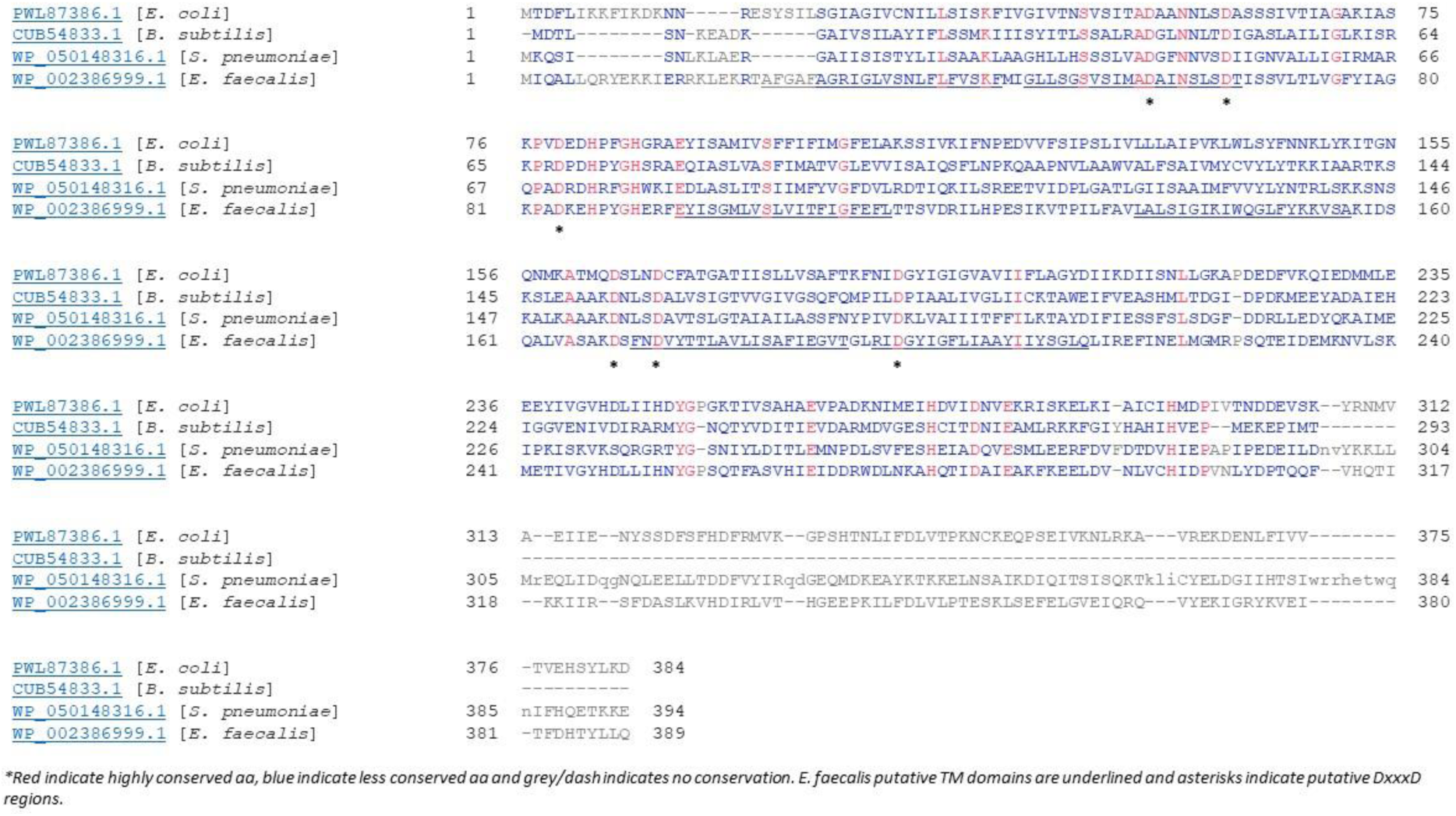
Sequence conservation of *E. faecalis* MntE.

**Figure S3.**
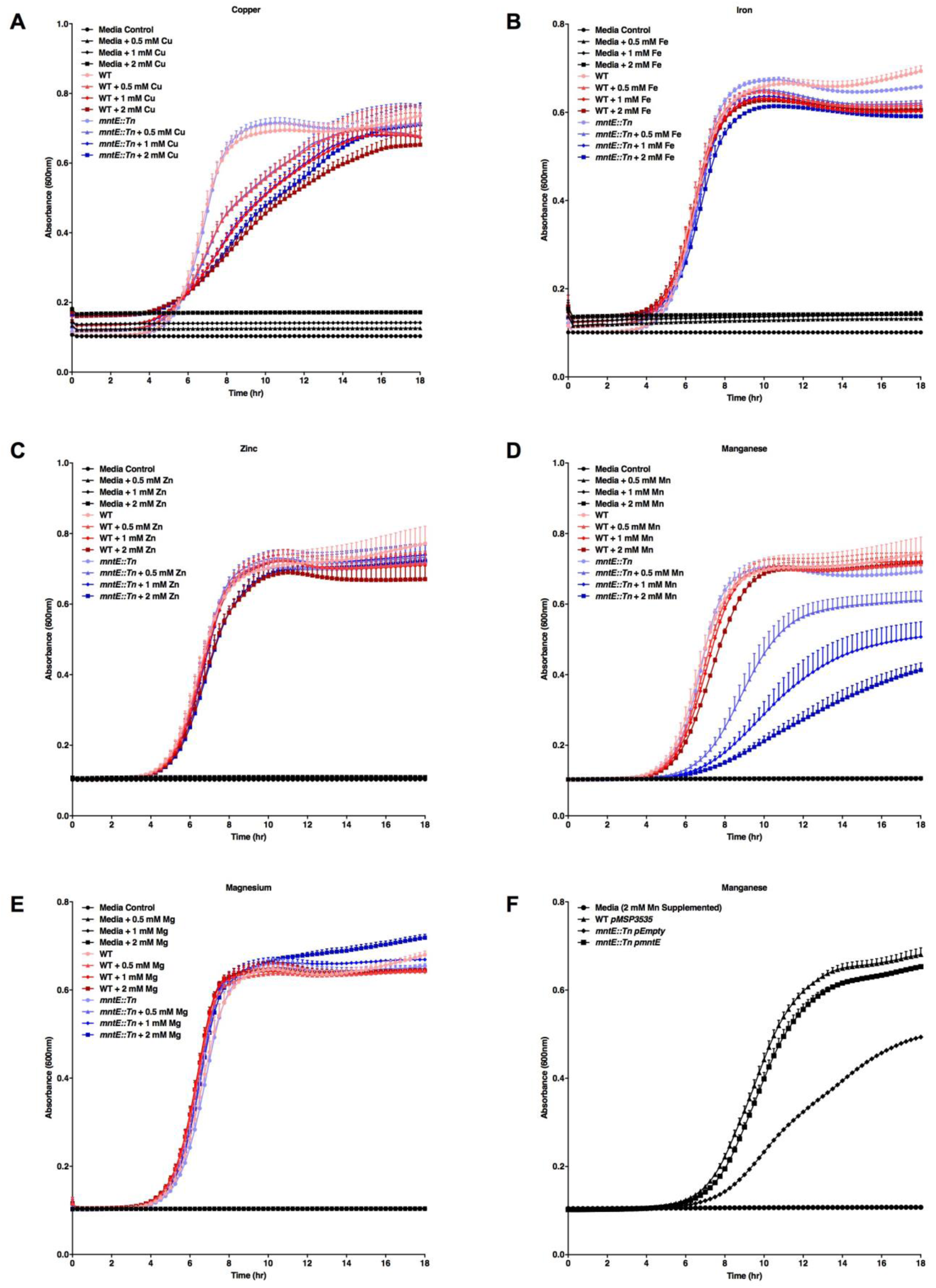
Cation sensitivity of MntE insertional deletion mutant. Planktonic growth of *E. faecalis* in TSBG and TSBG supplemented with increasing (A) copper, (B) iron, (C) zinc, (D) manganese, (E) magnesium and (F) manganese (complemented strain). Data points represent n =3 experiments, with three independent biological replicates averaged in each experiment.

**Figure S4.**
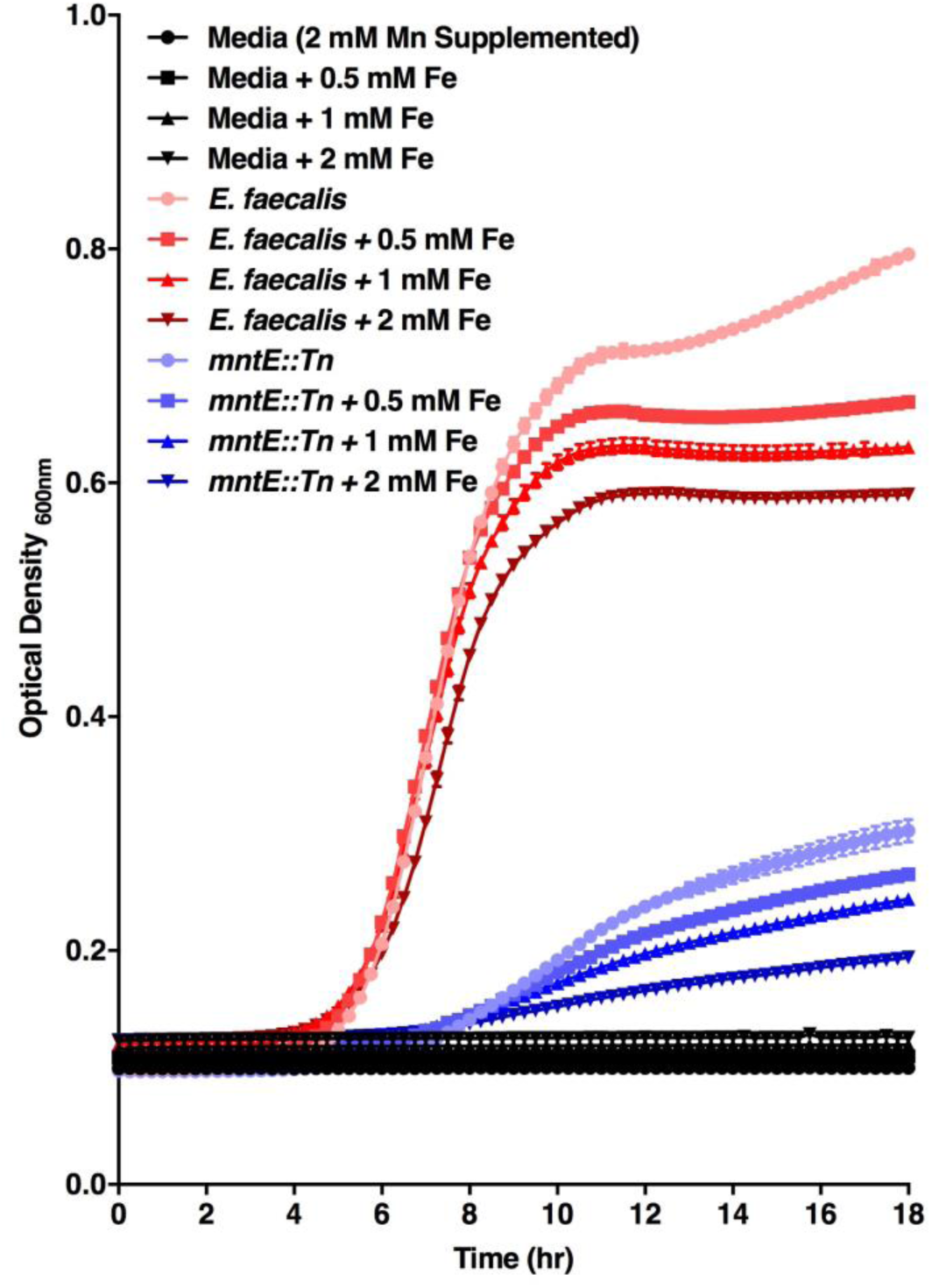
Fe supplementation does not alleviates Mn-mediated growth inhibition. Planktonic growth of *E. faecalis* in 2 mM Mn supplemented media with increasing iron concentration. Data points represent three independent biological replicates.

**Figure S5.**
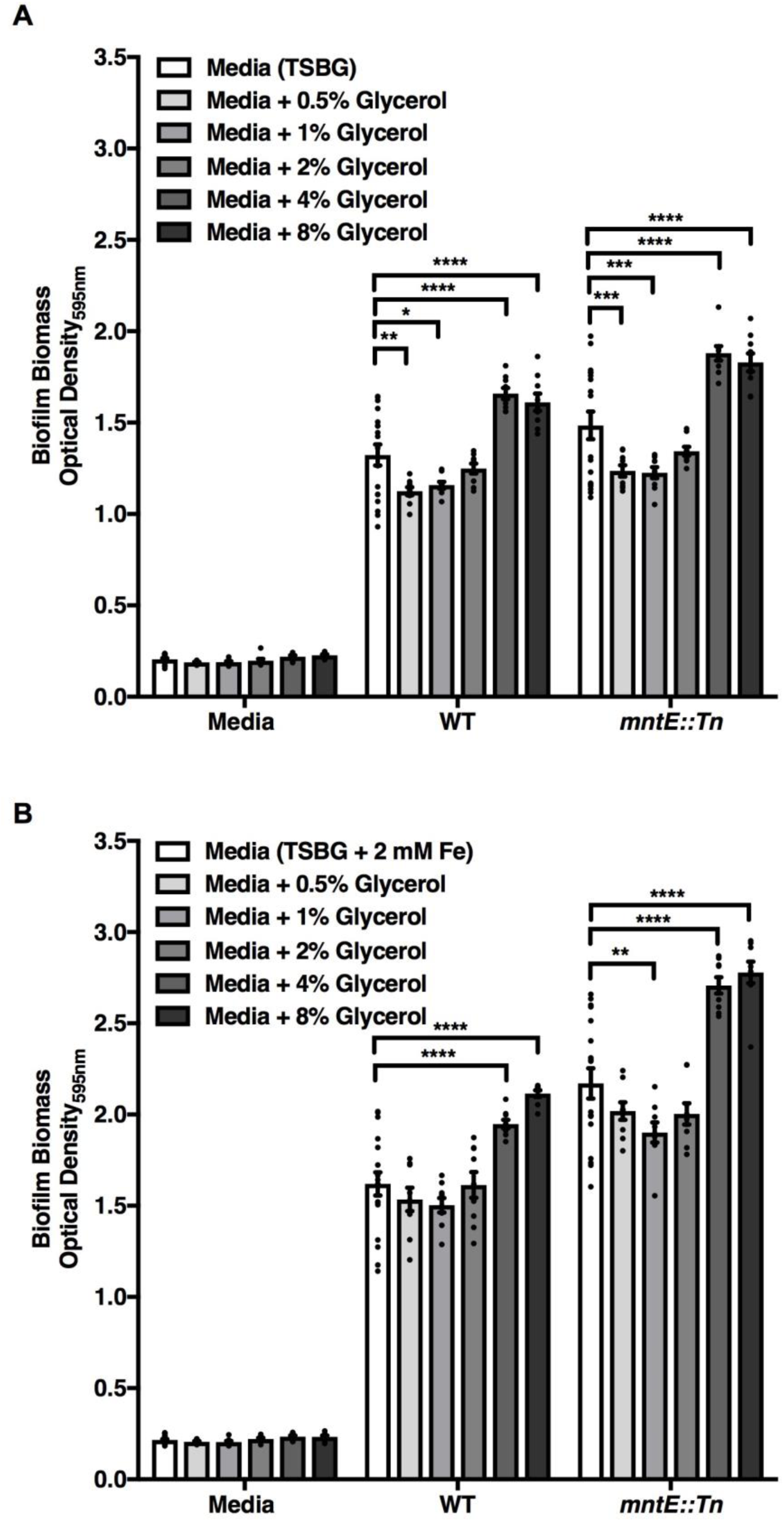
Glycerol supplementation promotes biofilm growth. Adherence biofilm biomass quantification of 120 hrs *E. faecalis* biofilm grown under glycerol supplementation in both (A) TSBG and (B) TSBG 2 mM Fe supplemented media. Data points represent at least six independent biological replicates assessed in at least n = 2 experiments. Statistical analysis was performed using two-way ANOVA with Bonferroni multiple comparison test. Error bar represents SEM.

**Figure S6.**
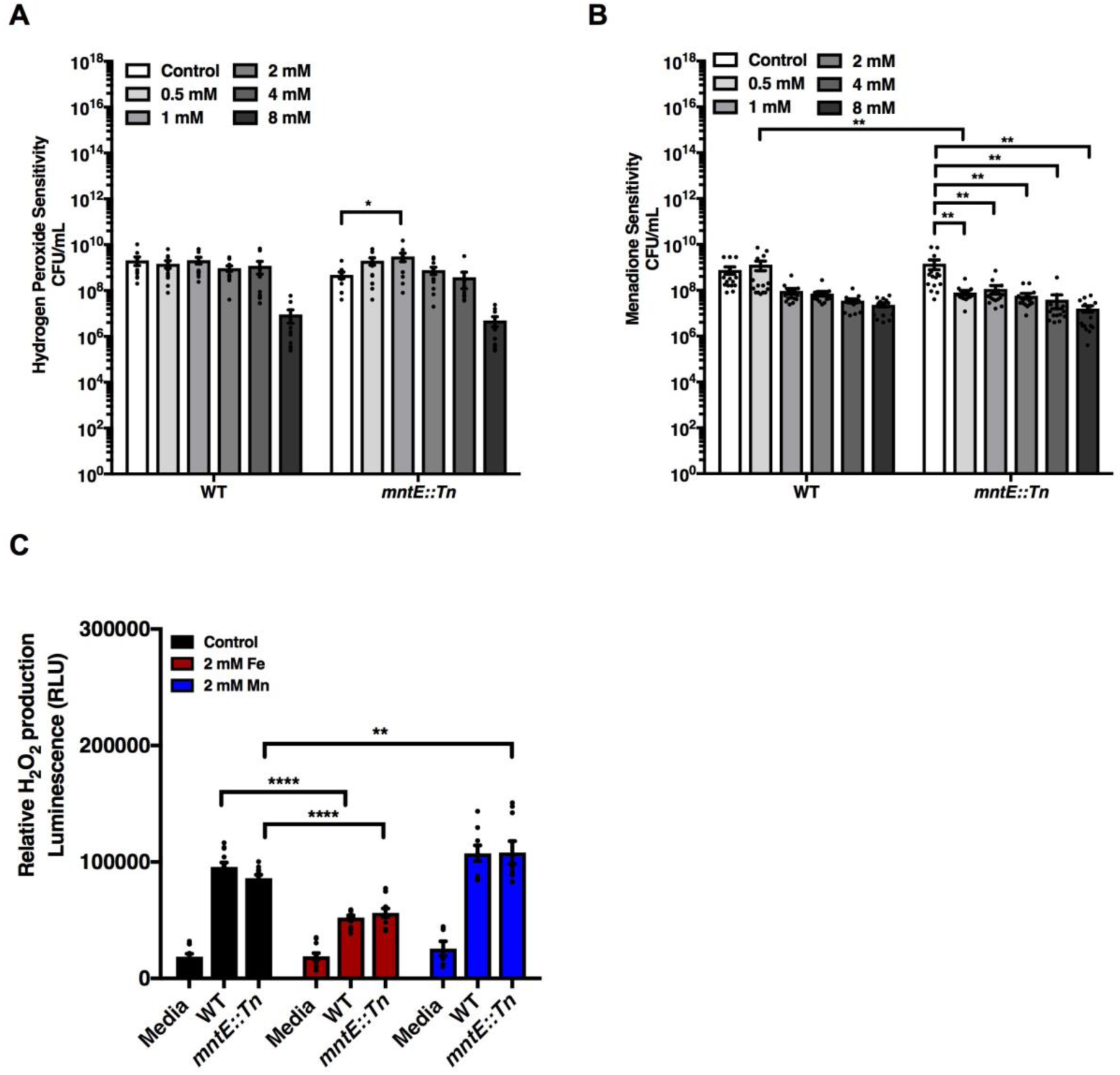
Absence of MntE does not impact oxidative stress tolerance and hydrogen peroxide production. CFU enumeration 2 hrs post exposure to (A) hydrogen peroxide and (B) menadione, and (C) hydrogen peroxide quantification based on arbitrary luminescence readings. Data points represent at least twelve independent biological replicates assessed in at least n = 4 experiments. Statistical analysis was performed using two-way ANOVA with Bonferroni multiple comparison test. Error bar represents SEM.

**Figure S7.**
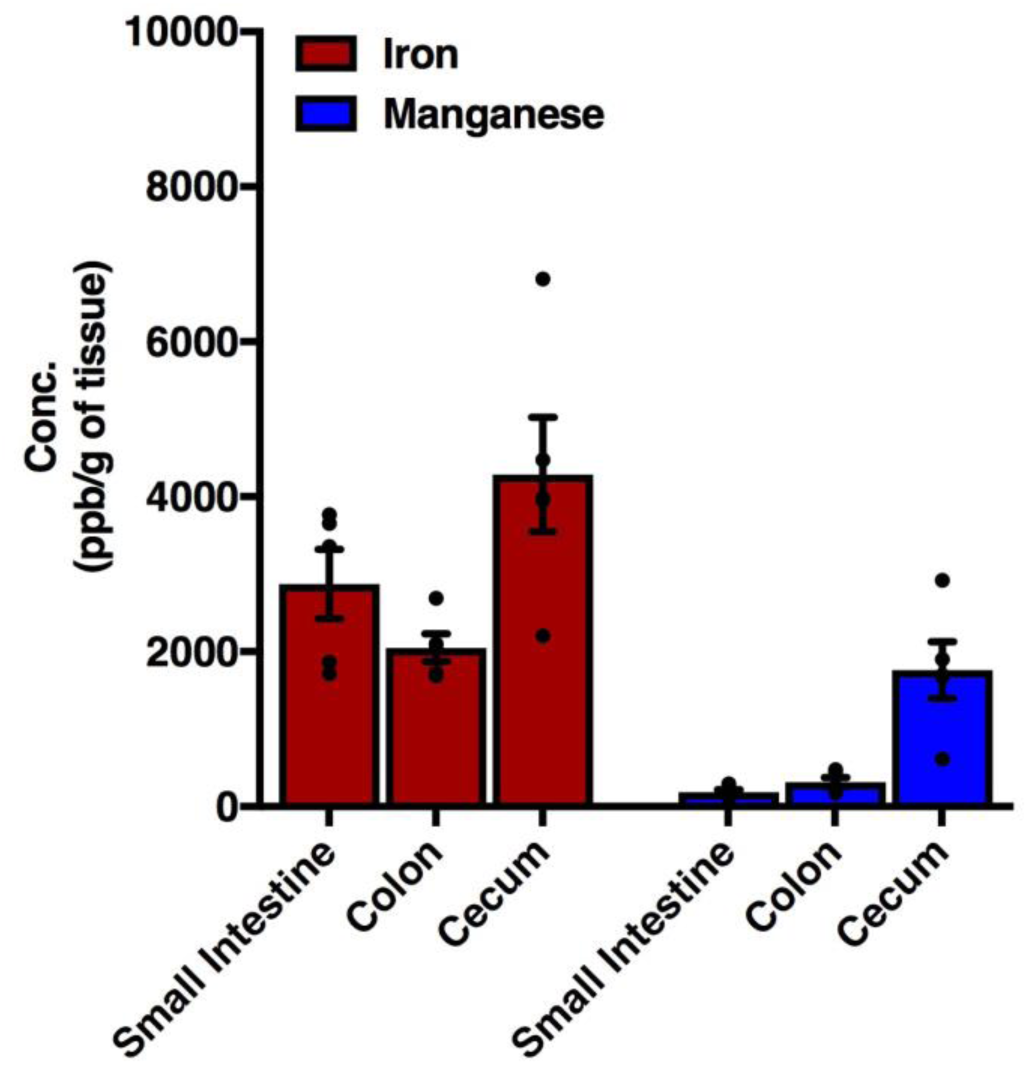
Iron and manganese abundance in mouse GI. ICP-MS analysis of iron and manganese levels from harvested GI tissues. Data points represent tissues harvested from five mice in one experiment.

**Supplementary Table S1.**
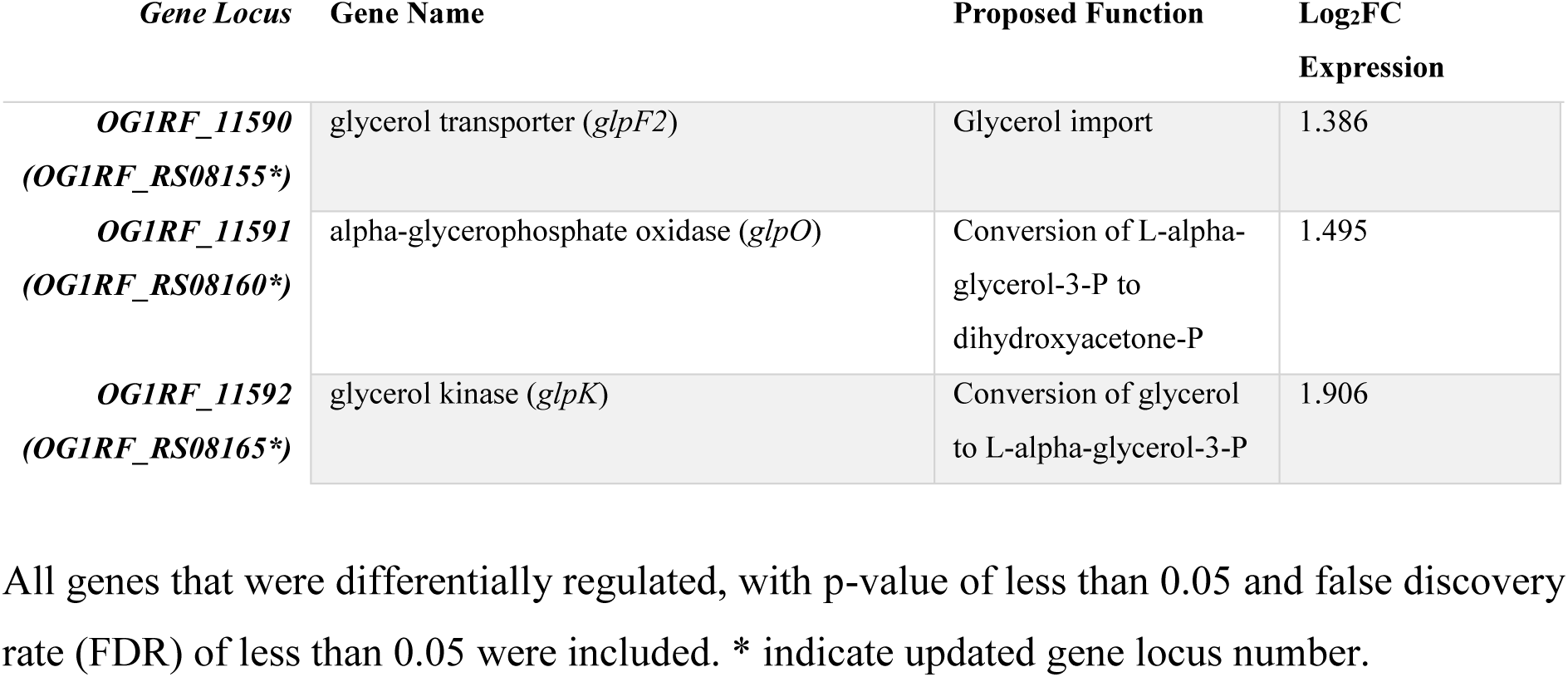
Global transcriptional changes in *mntE*∷Tn grown in Fe supplemented media compared with wild type.

| *Gene Locus* | *Gene Name* | *Proposed Function* | *Log<sub>2</sub>FC Expression* |
| --- | --- | --- | --- |
| ***OG1RF_11590***<br>**(*OG1RF_RS08155*\*)** | glycerol transporter (*glpF2*) | Glycerol import | 1.386 |
| ***OG1RF_11591***<br>**(*OG1RF_RS08160*\*)** | alpha-glycerophosphate oxidase (*glpO*) | Conversion of L-alpha-glycerol-3-P to dihydroxyacetone-P | 1.495 |
| ***OG1RF_11592***<br>**(*OG1RF_RS08165*\*)** | glycerol kinase (*glpK*) | Conversion of glycerol to L-alpha-glycerol-3-P | 1.906 |

All genes that were differentially regulated, with p-value of less than 0.05 and false discovery rate (FDR) of less than 0.05 were included. \* indicate updated gene locus number.

**Supplementary Table 2.**
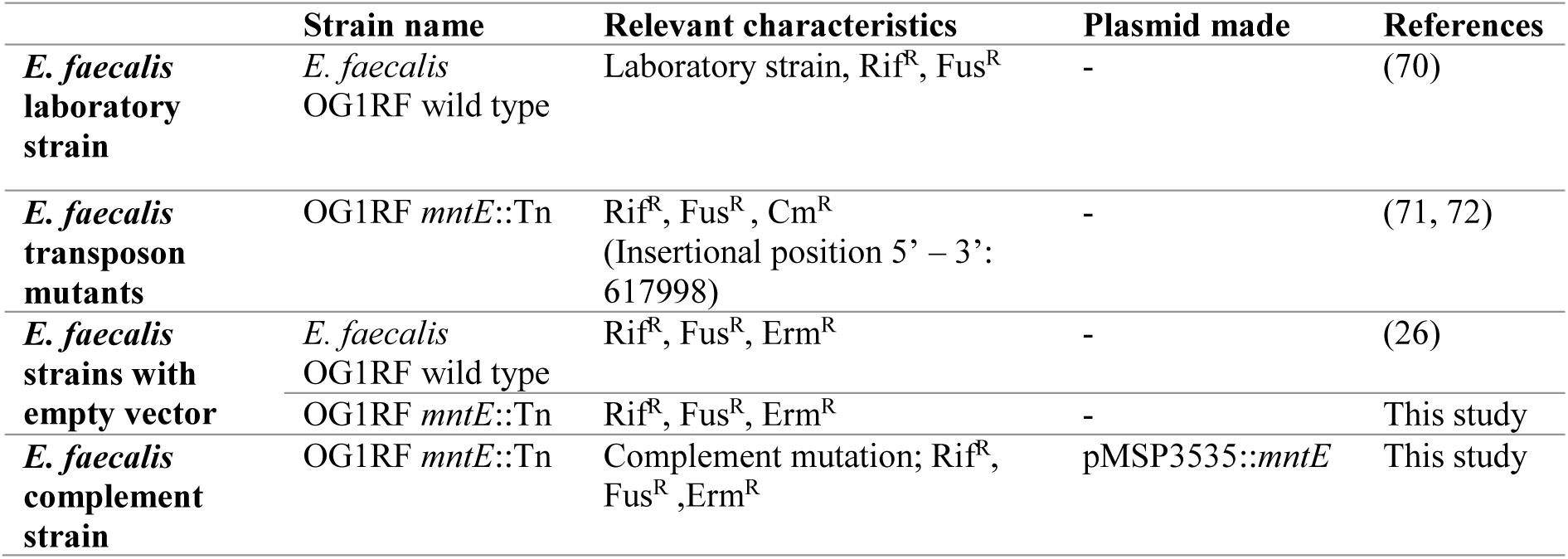
Strains and plasmids used in this study.

|  | **Strain name** | **Relevant characteristics** | **Plasmid made** | **References** |
| --- | --- | --- | --- | --- |
| ***E. faecalis* laboratory strain** | *E. faecalis* OG1RF wild type | Laboratory strain, Rif<sup>R</sup>, Fus<sup>R</sup> | - | (70) |
| ***E. faecalis* transposon mutants** | OG1RF *mntE*::Tn | Rif<sup>R</sup>, Fus<sup>R</sup>, Cm<sup>R</sup><br>(Insertional position 5' – 3': 617998) | - | (71, 72) |
| ***E. faecalis* strains with empty vector** | *E. faecalis* OG1RF wild type | Rif<sup>R</sup>, Fus<sup>R</sup>, Erm<sup>R</sup> | - | (26) |
|  | OG1RF *mntE*::Tn | Rif<sup>R</sup>, Fus<sup>R</sup>, Erm<sup>R</sup> | - | This study |
| ***E. faecalis* complement strain** | OG1RF *mntE*::Tn | Complement mutation; Rif<sup>R</sup>, Fus<sup>R</sup>, Erm<sup>R</sup> | pMSP3535::*mntE* | This study |

**Supplementary Table 3.**
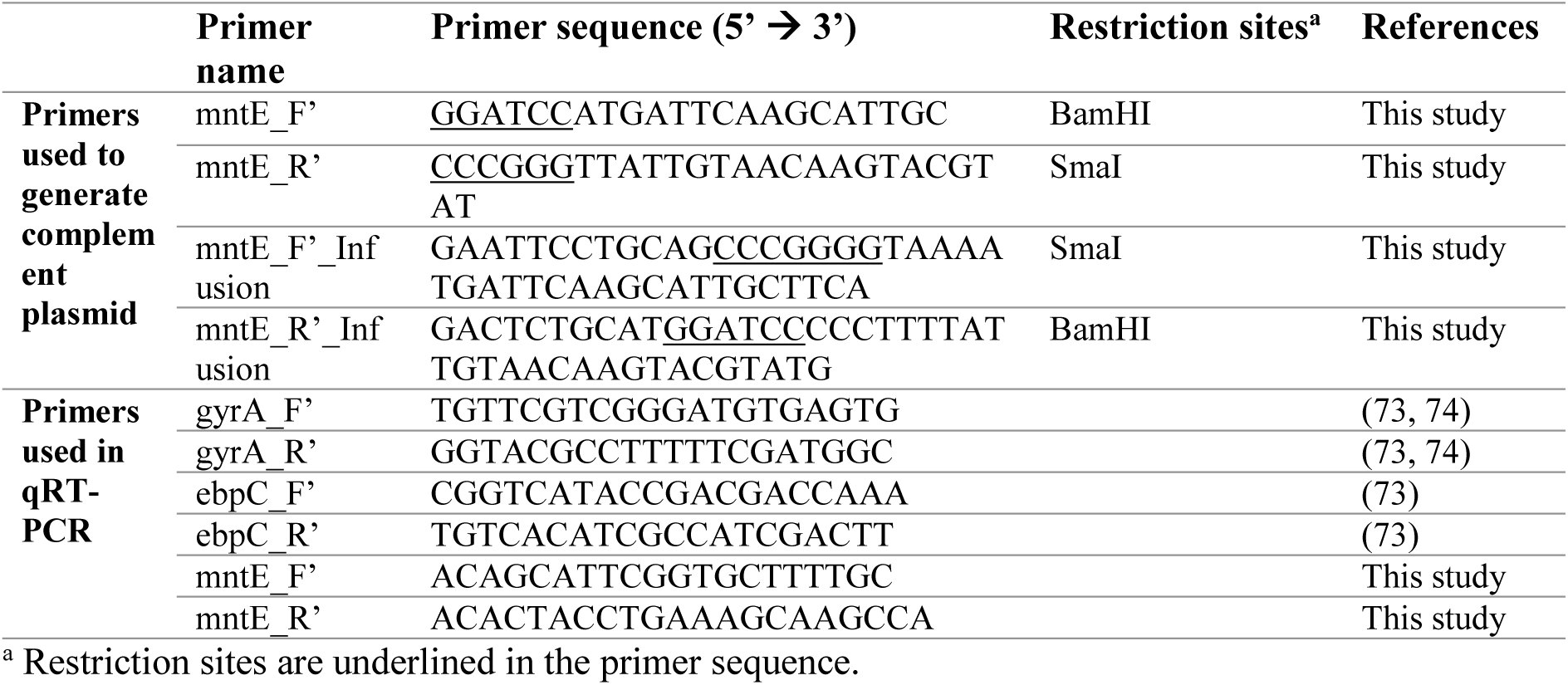
Primers used in this study.

